# IsdH binding to hemoglobin:haptoglobin is decoupled from iron acquisition in *Staphylococcus aureus*

**DOI:** 10.64898/2026.06.17.732818

**Authors:** Valeria Buoli Comani, Omar De Bei, Sarah Hijazi, Antonio Cataldi, Giulia Paris, Giulia Sassi, Marialaura Marchetti, Barbara Campanini, Luca Ronda, Serena Faggiano, Ben F. Luisi, Emanuela Frangipani, Stefano Bettati

**Affiliations:** Department of Food and Drug, University of Parma, Parco Area delle Scienze, 27/A, Parma, Italy; Interdepartmental Center Biopharmanet-TEC, University of Parma, Tecnopolo Padiglione 33, Parma, Italy; Department of Medicine and Surgery, University of Parma, Via Gramsci, 14, Parma, Italy; Department of Biomolecular Sciences, University of Urbino Carlo Bo, Via Santa Chiara, 27, Urbino, Italy; Department of Biochemistry, Sanger Building, University of Cambridge, Cambridge, UK; Department of Chemistry, Life Sciences and Environmental Sustainability, University of Parma, Parco Area delle Scienze, 27/A, Parma, Italy; Institute of Biophysics, CNR, Via G. Moruzzi, 1, Pisa, Italy

**Keywords:** Staphylococcus aureus, Hemoglobin, Haptoglobin, IsdH, Iron acquisition

## Abstract

*Staphylococcus aureus* requires iron for proliferation during infection and acquires it mainly from host hemoglobin (Hb) through the iron-regulated surface determinant (Isd) system. The hemophores IsdB and IsdH mediate the initial steps of Hb recognition. Notably, IsdH also recognizes the hemoglobin:haptoglobin (HbHp) complex, although the molecular determinants and physiological relevance of this interaction remain unclear. Here, combining cryo-electron microscopy and biochemical and cellular assays, we define the basis of HbHp recognition by IsdH. The structure reveals how IsdH engages HbHp and captures the intrinsic conformational flexibility of the HbHp assembly. Functional analyses demonstrate that, despite retaining the ability to extract heme from HbHp, IsdH does not support *S. aureus* growth under iron-restricted conditions when HbHp is the sole iron source. These findings suggest that HbHp recognition by IsdH may serve functions beyond nutrient iron acquisition, contributing to modulation of host-pathogen interactions.

## Introduction

*Staphylococcus aureus* is a major human pathogen responsible for a wide spectrum of infections that include bacteremia, endocarditis, and deep tissue infections (Tong *et al*, 2015; Wertheim *et al*, 2005). Successful colonization and proliferation critically depend on the ability of the bacterium to acquire essential nutrients from the host environment (Balasubramanian *et al*, 2017). Among these, iron is indispensable yet scarce due to host-imposed sequestration mechanisms collectively referred to as Nutritional Immunity (NI) (Hood & Skaar, 2012). During infection, pathogens encounter an environment depleted of accessible iron, forcing them to exploit host-associated iron reservoirs (Marchetti *et al*, 2020), such as the heme prosthetic group bound to hemoglobin (Hb) inside erythrocytes (Vogt *et al*, 2021). Upon hemolysis, extracellular Hb is captured by the plasma glycoprotein haptoglobin (Hp), forming a stable HbHp complex that prevents heme-mediated oxidative damage (Andersen *et al*, 2017) (**Figure 1a**). Hp is a hetero-oligomeric protein built from αβ protomers, in which the α-chains mediate oligomerization, whereas the β-chains mediate Hb binding (Mollan *et al*, 2014; Wejman *et al*, 1984). The HbHp complexes are efficiently recognized and cleared by the scavenger receptor CD163 expressed on macrophages and monocytes (Zhou & Higgins, 2025; Etzerodt *et al*, 2024; Xu *et al*, 2025; Huang *et al*, 2025), thereby limiting heme availability and reinforcing host NI (**Figure 1a**).

**Figure 1.**
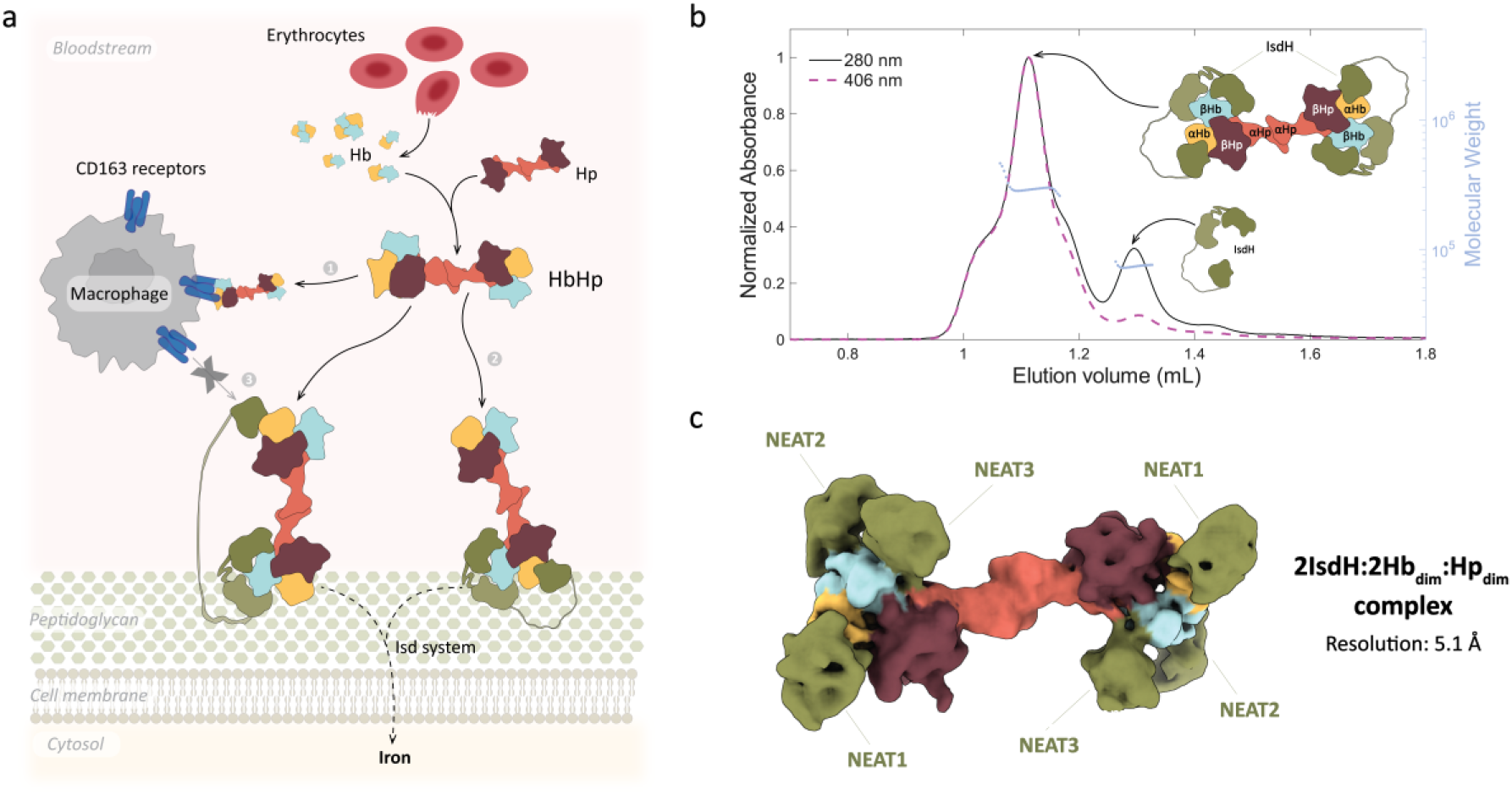
IsdH:HbHp complex. (**a**) Schematic representation of the IsdH-mediated hijacking of the HbHp complexes by *Staphylococcus aureus*. Following erythrocyte hemolysis, extracellular hemoglobin (Hb) is released into the bloodstream and dissociates from its tetrameric form into αβ dimers, which are rapidly scavenged by haptoglobin (Hp). Under physiological conditions, HbHp complexes are cleared from circulation through recognition by the macrophage receptor CD163 (1). *S. aureus* exploits this host pathway through the hemophore IsdH, which binds HbHp complexes and is proposed to promote heme extraction and subsequent internalization as an iron source (2). IsdH binding may additionally interfere with the interaction between HbHp complexes and CD163, thereby perturbing physiological HbHp clearance (3). (**b**) Size-exclusion chromatography coupled to light scattering (SEC–LS) of a mixture of Hp, metHb and IsdH. Elution was monitored at 280 nm (aromatic residues, solid black) and 406 nm (heme, dashed magenta). Main peaks correspond to species represented by the icons: 2IsdH:2Hb_dim_:Hp_dim_ and free IsdH. (**c**) Cryo-EM reconstruction of the fully assembled ternary complex 2IsdH:2Hb_dim_:Hp_dim_ at 5.1 Å (EMD-57414).

To overcome this restriction, *S. aureus* has evolved the iron-regulated surface determinant (Isd) system, a coordinated pathway that enables the capture of Hb and the extraction and transport of heme across the cell envelope (Mazmanian *et al*, 2003). The initial step of this process is mediated by surface-anchored hemophores containing NEAr iron Transporter (NEAT, N) domains, which recognize Hb and extract heme; among these, IsdB and IsdH, composed of two and three NEAT domains respectively, act as primary Hb receptors.

Despite their apparent functional overlap, these receptors are not fully redundant (Buoli Comani *et al*, 2025). IsdB is highly efficient at extracting heme from Hb (Bowden *et al*, 2018; Gianquinto *et al*, 2019; De Bei *et al*, 2022, 2025), while IsdH, originally identified as Haptoglobin receptor A (HarA), was discovered as capable of recognizing both free Hb and HbHp complexes (Dryla *et al*, 2003). Subsequent functional investigations have largely relied on minimal functional constructs, in particular the NEAT2-Linker2-NEAT3 region (IsdH^N2N3^), competent for Hb binding and heme extraction (Buoli Comani *et al*, 2025; Dryla *et al*, 2007; Krishna Kumar *et al*, 2011; Pilpa *et al*, 2009; Sjodt *et al*, 2016; Spirig *et al*, 2013; Valenciano-Bellido *et al*, 2022). Within this framework, Mikkelsen et al. (Mikkelsen *et al*, 2020) showed that the presence of Hp can inhibit IsdH-mediated heme extraction. In turn, IsdH might also restrict access to the HpHb complex by the macrophage-associated CD163 receptors (Sæderup *et al*, 2016)(**Figure 1a**). However, these studies are inherently limited by the use of truncated constructs and by mutations that impair heme extraction, thus providing only a partial view of the interaction (*e*.*g*., PDB: 6TB2).

Recent analyses of IsdH have shown that the protein is unexpectedly inefficient in heme extraction from free Hb under physiological timescales in comparison with IsdB (Comani *et al*, 2026). On the other hand, IsdH binds the HbHp complex and directly interferes with host mechanisms for Hb clearance (Sæderup *et al*, 2016).

Beyond *S. aureus*, only a limited number of pathogens have evolved the ability to exploit the HbHp complex as a heme source. Gram-negative *Neisseria* species use the TonB-dependent receptor HpuAB to bind and internalize HbHp (Rohde & Dyer, 2004; Chen *et al*, 1996), while hematophagous protozoa such as *Trypanosoma brucei* rely on HpHbR to endocytose the complex (Higgins *et al*, 2013). Additional examples include outer-membrane proteins in *Haemophilus influenzae* (Seale *et al*, 2006) and surface receptors in *Corynebacterium diphtheriae* (Lyman *et al*, 2021), which support HbHp recognition and utilization, highlighting the rare but convergent strategies microbes have evolved to access heme from this host defense complex.

A coherent mechanistic understanding of HbHp recognition and its role in bacterial proliferation by staphylococcal IsdH is still lacking. In particular, it remains unclear how full-length IsdH engages the HbHp complex at the molecular level, whether this interaction supports efficient heme acquisition under iron-limiting conditions, and to what extent it underpins the functional specialization of IsdH compared to IsdB.

By combining cryo-electron microscopy (cryo-EM) with biochemical and cellular assays, here we investigate the structural and functional interplay between IsdH and the HbHp complex. Although IsdH is capable of extracting heme from HbHp, our data show that this interaction does not support *S. aureus* growth under iron-restricted conditions. These findings indicate that heme extraction from HbHp is not sufficient to sustain bacterial proliferation and point to a distinct role for IsdH at the host-pathogen interface.

## Results

### IsdH Binding to the Hemoglobin-Haptoglobin Complex

To define the oligomeric state and stoichiometry of the assembly used for structural analysis, we first characterized the interaction between IsdH, Hb, and Hp in solution. We employed metHb, the oxidized form of Hb that preferentially associates with Hp *in vivo*, together with the dimeric Hp1-1 (Wejman *et al*, 1984), which avoids the miscellaneous oligomerization typical of other Hp variants and provides a homogeneous system suitable for cryo-EM analysis (Zhou & Higgins, 2025; Etzerodt *et al*, 2024; Xu *et al*, 2025; Huang *et al*, 2025, 163).

Size-exclusion chromatography coupled to light scattering detection (SEC-LS) was used to characterize the individual components and complex formation upon mixing Hp, metHb, and IsdH (**Figure 1b, Figure S1, Table S1**). Consistent with previous reports (Andersen *et al*, 2012), HbHp eluted as a monodisperse species corresponding to an Hp dimer bound to two Hb dimers (2Hb_dim_:Hp_dim_; **Figure S1d, Table S1**). Upon addition of IsdH, the elution profile shifted toward a higher apparent molecular weight, and the main peak corresponded to a species compatible with a ternary complex composed of two IsdH molecules bound to HbHp (2IsdH:2Hb_dim_:Hp_dim_; **Figure 1b** and **Figure S1h**). A second well-defined peak eluting at higher elution volumes was also observed and corresponded to free IsdH in solution (**Table S1**). Notably, further increasing the concentration of IsdH did not lead to the formation of IsdH:HbHp assemblies containing additional IsdH molecules (**Figure S1f, Table S1**).

Cryo-EM was performed in the same condions previously characterized by SEC-LS. Single-particle analysis (SPA) yielded a three-dimensional reconstruction of the IsdH:HbHp complex at an overall resolution of 5.1 Å (EMD-57414; **Figure 1c** and **Figure S2**). The reconstruction revealed the presence of two IsdH molecules bound to the HbHp complex, in agreement with the stoichiometry identified by SEC-LS. Inspection of the cryo-EM micrographs also revealed the presence of low-order aggregates in the preparation (**Figure S3a**), which are consistent with the shoulder eluting before the main SEC-LS peak. More interestingly, SPA identified a second population of particles that yielded a reconstruction of a smaller assembly at 3.8 Å resolution (EMD-57413, **Figure S3b**). The structure revealed an Hp dimer bound to a single Hb dimer and one IsdH molecule, defining a 1IsdH:1Hb_dim_:Hp_dim_ assembly. Because IsdH specifically recognizes Hb rather than Hp alone, the absence of the second Hb dimer implies that only a single IsdH molecule can be accommodated in this assembly. Accordingly, this population likely corresponds to the shoulder eluting at slightly higher volumes than the main peak in the SEC-LS chromatogram. This assignment is further supported by the chromatographic absorbance profile (**Figure 1b**), in which the relative intensity of the heme signal at 406 nm is reduced compared to the protein signal at 280 nm, consistent with a species containing fewer heme groups than the fully assembled ternary complex.

### Flexibility analysis of the 2IsdH:2Hb_dim_:Hp_dim_ complex

The cryo-EM reconstruction of the 2IsdH:2Hb_dim_:Hp_dim_ assembly (≈290 kDa) did not exceed ∼5 Å resolution, suggesting that conformational heterogeneity might limit map refinement. To investigate this possibility, we applied two complementary approaches for flexibility analysis: Multi-body Refinement (**Figure 2a**, RELION pipeline (Scheres, 2012)) and 3DFlex (**Figure 2b**, cryoSPARC pipeline (Punjani & Fleet, 2023)) (https://figshare.com/s/e0818ca4ffbaf596da81). Multi-body Refinement models conformational variability by defining rigid regions that move relative to one another. Based on the architecture of the complex, we defined bodies corresponding to the two 1IsdH:1Hb_dim_:Hp_mon_ subunits (**Figure 2c**). In contrast, 3DFlex does not rely on predefined masks and learns continuous structural variability directly from the particle dataset, yielding a refined reconstruction with improved local resolution (**Figure 2d**).

**Figure 2.**
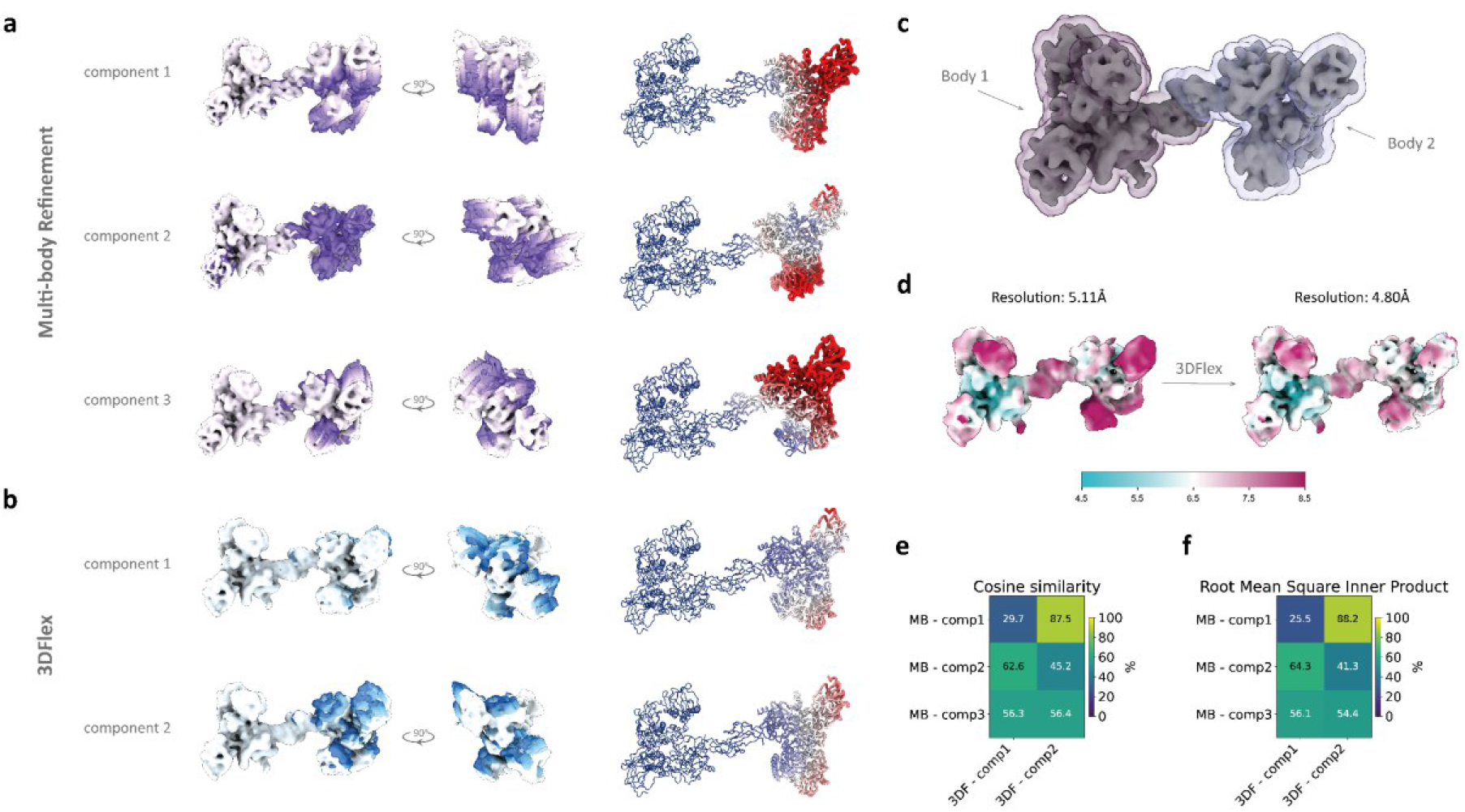
Flexibility analysis of the 2IsdH:2Hb_dim_:Hp_dim_ complex. (**a**) Principal components of the motion identified by multi-body refinement. For each component, reconstructed maps representing the motion are shown from two orthogonal views. Conformational variability is also visualized on the models by scaling backbone thickness and color according to the computed B-factors (blue, low mobility; red, high mobility). (**b**) Principal components of the motion identified by 3DFlex. As in (**a**), maps corresponding to each motion component are shown from two views and the model displays mobility through backbone thickness and B-factor–based color scaling. (**c**) Definition of rigid bodies used for Multi-body Refinement. (**d**) Local resolution maps of the 2IsdH:2Hb_dim_:Hp_dim_ complex before and after 3DFlex. (**e**,**f**) Comparison between motion components identified by the Multi-body Refinement (MB) and 3DFlex (3DF) using (**e**) cosine similarity and (**f**) Root Mean Square Inner Product (RMSIP) analyses.

Both methods decompose conformational variability into principal components describing dominant motions within the dataset. Multi-body Refinement identified three principal components, whereas 3DFlex identified two. To visualize these motions, atomic models were flexibly fitted into reconstructed maps using IMOD (https://bio3d.colorado.edu/imod/), and the resulting trajectories were represented on the models by scaling backbone thickness and color according to the computed B-factor, ranging from blue (low mobility) to red (high mobility) (**Figure 2a,b**). To determine whether the motions identified by the two approaches corresponded to similar structural transitions, we compared their principal components using cosine similarity (**Figure 2e**) and root mean square inner product (RMSIP) analyses (**Figure 2f**).

Comparable motions were detected both with and without predefined body masks (*i*.*e*. the first 3DFlex component corresponds to the second and third Multi-body Refinement components, whereas the second 3DFlex component mainly matches the first one), indicating that the observed flexibility reflects an intrinsic movement of the complex. In particular, the hinge region underlying this motion can be localized at the interface between the Hp α-chains, suggesting that it mediates the relative displacement of the two halves of the complex.

These results, viewed in a broader context, are consistent with the only available cryo-EM data on HbHp assemblies, in which pronounced structural flexibility was also observed. In particular, recent cryo-EM studies of the HbHp complex bound to the CD163 receptor resolved only part of the HbHp assembly (Zhou & Higgins, 2025; Etzerodt *et al*, 2024; Xu *et al*, 2025; Huang *et al*, 2025), supporting the presence of substantial inter-domain mobility.

### IsdH:HbHp complex: The role of Linker1

Analysis of the intrinsic flexibility of the assembly enabled identification and discretization of common motions within the complex but did not improve the resolution of the maps. The best reconstruction was achieved by focusing on the minimal stable unit of the assembly. A focused refinement on half of the assembly (1IsdH:1Hb_dim_:Hp_mon_) yielded the best reconstruction, reaching a final resolution of 3.3 Å (EMD-57212, PDB: 29JK, **Figure 3a and Figure S4a**). In this reconstruction, the NEAT2-NEAT3 region of IsdH is bound to the β-chain of Hb and captures the heme in the extracted state, coordinated by NEAT3 (**Figure S4b**). The corresponding β-subunit therefore adopts an apo configuration and, consistent with previous reports, the F helix lacks interpretable density (Bowden *et al*, 2018; De Bei *et al*, 2022; Ellis-Guardiola *et al*, 2020) (**Figure S4d**). The NEAT1 domain engages the α-chains of Hb within the complex.

**Figure 3.**
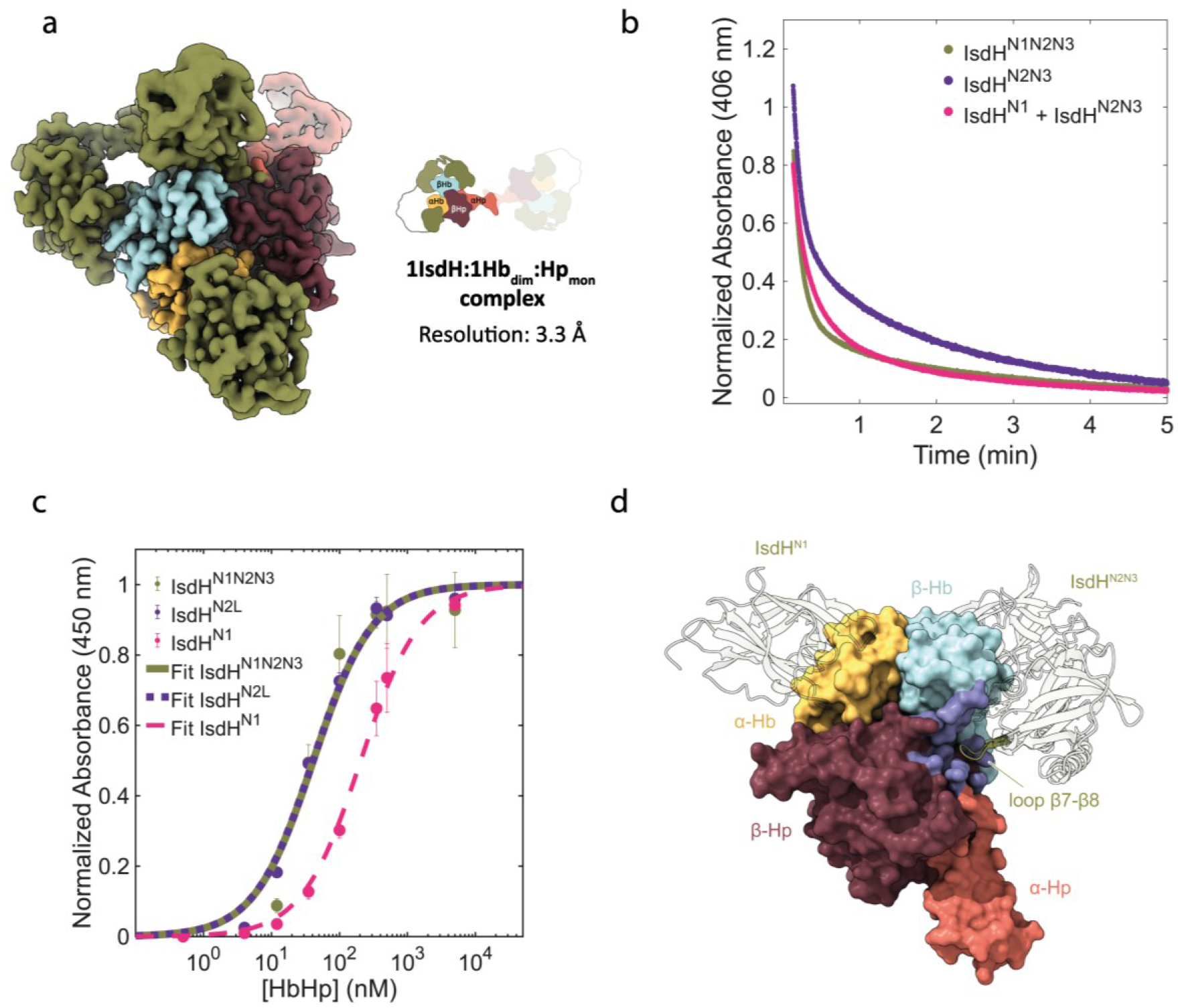
Structural and functional analysis of IsdH:HbHp. (**a**) Cryo-EM map of the 1IsdH:1Hb_dim_:Hp_mon_ complex reconstructed at 3.3 Å resolution (EMD-57212). (**b**) Kinetics of heme extraction from the metHb:Hp1–1 complex by IsdH^N1N2N3^ (green), IsdH^N2N3^ (purple), and IsdH^N2N3^ in the presence of isolated IsdH^N1^ (magenta), monitored by visible spectroscopy at 406 nm in PBS (pH 7.4). (**c**) ELISA-based estimation of the affinity of IsdH constructs for the HbHp complex. Measured apparent dissociation constants (K_D_) are: 41 ± 27 nM for IsdH^N1N2N3^; 41 ± 13 nM for IsdH^N2L^; 206 ± 40 nM for IsdH^N1^. (**d**) Surface representation of the HbHp complex colored as in panel **a**. IsdH is shown as a semi-transparent cartoon. The NEAT3 β7-β8 loop (residues 636-644) is highlighted in green and inserts into a cleft formed at the HbHp interface (purple).

A dense network of intermolecular contacts was identified using RING (**Figure S4**), showing that NEAT1 interactions with the α-chains are highly consistent with those reported by crystallography (PDB: 3SZK; **Table S3**). Likewise, most contacts observed for the NEAT2-NEAT3 region bound to the β-chains are comparable to those reported for the same domains interacting with the α-chains of Hb in the only available cryo-EM structure of full-length IsdH in complex with free Hb, obtained in the absence of Hp (Comani *et al*, 2026) (PDB: 9SZW; **Table S4**).

The ∼100-residue segment connecting NEAT1 to NEAT2 (Linker1), however, lacks interpretable density in the cryo-EM map, indicating pronounced conformational flexibility. AlphaFold3 predicts this region to be unfolded (**Figure S5 a**,**b**), and sequence-based analyses classify it as an entropic chain containing potential protein-protein interaction motifs (**Figure S5 c**,**d**). If Linker1 formed stable interactions with the HbHp complex, it would be expected to adopt an ordered conformation upon binding. However, the absence of density indicates that the region remains highly flexible and does not participate in stable contacts within the resolved assembly.

As the structural analysis did not reveal a defined role for Linker1, biochemical and biophysical approaches were employed to probe its functional relevance. Heme extraction assays indicated that Linker1 is not strictly required for IsdH heme scavenging activity (**Figure 3b**). The IsdH^N2N3^ construct displays reduced extraction efficiency relative to the full-length protein. However, adding the isolated NEAT1 domain (*i*.*e*., without Linker1) *in trans* restores activity to levels comparable to the full-length construct. This result demonstrates that NEAT1 can promote heme extraction even when not covalently linked to the rest of the protein. Binding measurements further showed that the affinity of IsdH for HbHp complex was comparable to that of the IsdH^N2L^ construct, while NEAT1 alone displayed substantially lower affinity (**Figure 3c**). Given the structural organization of the IsdH:HbHp complex – where NEAT1 and the NEAT2-NEAT3 region engage Hb α- and β-chains, respectively – and the absence of steric constraints on NEAT2-NEAT3 binding to the α-chains in the HbHp complex (Mikkelsen *et al*, 2020) (PDB: 6TB2), the relative affinities measured here would in principle suggest that NEAT2-NEAT3 could outcompete NEAT1 for α-chain binding. However, this scenario is not supported by our data, as both the cryo-EM reconstruction (**Figure S3b**) and SEC-LS analysis (**Figure S1f**) consistently indicate binding of a single IsdH molecule per Hb dimer. In agreement with these observations, heme extraction from HbHp appears incomplete compared with the extraction experiment performed using Hb alone (**Figure S6**).

This behavior suggests that structural features of the complex limit the access of additional IsdH molecules. Linker1 does not directly contribute to ligand recognition but may instead play a structural or steric role. In this framework, the flexible linker may act as a dynamic envelope around the IsdH:HbHp assembly that restricts access of additional IsdH molecules in the *in vitro* assay. Such a mechanism may be particularly relevant *in vivo*, where IsdH is anchored to the bacterial cell wall, and uncontrolled multivalent interactions with surrounding proteins could otherwise occur.

### Hp redirects IsdH binding selectivity, favoring a more persistent complex

The structural assembly presented here (PDB: 29JK), together with the full-length IsdH:Hb complex (Comani *et al*, 2026)(PDB: 9SZW), provides a framework to compare the domain organization and positioning of IsdH at the initial encounter with Hb, either in complex with Hp or in its free form. IsdH contains two domains specialized in Hb binding (*i*.*e*., NEAT1 and NEAT2), while only NEAT3 is responsible for heme extraction. Functional interest has long converged on the NEAT2-NEAT3 region, which has been identified as the minimal functional construct required for efficient heme acquisition (Sjodt *et al*, 2016; Spirig *et al*, 2013; Valenciano-Bellido *et al*, 2022). Interestingly, the presence of Hp shifts the chain selectivity of this region from the α-chains in free Hb to the β-chains in the HbHp complex.

Structural inspection identifies a single IsdH region directly influenced by Hp binding that can redirect NEAT2-NEAT3 from the α-to the β-chain: the NEAT3 β7-β8 loop (residues 636–644, **Figure 3d**). In the absence of Hp, this loop contacts Hb, burying ∼110 Å^2^ of surface area. In the HbHp complex, however, Hb and Hp together form a composite cleft that accommodates the loop, increasing the buried surface area to ∼191 Å^2^. Across all available IsdH structures, this loop represents the most conformationally dynamic region of the Hb receptor (**Figure S7 a-c**), consistent with Hydrogen-Deuterium Exchange Mass Spectrometry (HDX–MS)(Valenciano-Bellido *et al*, 2022) and molecular dynamics analyses (Ellis-Guardiola *et al*, 2020), while functional studies identify several residues within this region as critical determinants of productive heme extraction (**Figure S7d**).

Notably, both α- and β-chains of Hb present accessible binding surfaces that can accommodate the NEAT3 β7-β8 loop. Consistently, in the IsdH:Hb complex (Comani *et al*, 2026) (PDB: 9SZW), the loop adopts a properly positioned and folded conformation even in the absence of Hp. However, the composite cleft described above emerges only upon HbHp formation. These observations indicate that, although NEAT3 can engage both α- and β-chains, its contribution to chain selectivity becomes functionally relevant only in the presence of Hp, where the composite interface stabilizes and biases the encounter complex. In its absence, as in the IsdH:Hb complex where the NEAT2-NEAT3 region is bound to α-chains, NEAT3 does not appear to dictate the initial chain preference. Instead, early selectivity is likely governed by NEAT2, consistent with truncated constructs lacking functional NEAT3 (*e*.*g*., IsdH^N2L^) which preferentially associate with α-chains in solution (Buoli Comani *et al*, 2025). By contrast, engagement of the β7-β8 loop within the HbHp interface enables NEAT3 to contribute to early recognition, biasing engagement toward β-chains.

In addition to redirecting chain selectivity, the HbHp composite cleft appears to exert a pronounced stabilizing effect on the whole NEAT3 domain. Indeed, although this domain has previously been described as highly flexible and poorly resolved in cryo-EM reconstructions of IsdH:Hb complexes (Buoli Comani *et al*, 2025), it appears comparatively well ordered in the present structure despite having already acquired heme. Consistently, previously reported structural data of the destabilizing Y642A mutant (**Figure S7 e-f**) show that the β7-β8 loop remains ordered only when engaged within the HbHp composite cleft. Notably, such stabilization may not necessarily favor efficient downstream heme transfer, since persistent association with Hb could reduce accessibility to other components of the Isd system. In parallel, as discussed above, the N-terminal N1–Linker1 region also appears to contribute primarily to stabilization of the complex with HbHp rather than to maximizing extraction efficiency. Altogether, these findings support the idea that IsdH may have principally evolved to maintain a persistent interaction with Hb when the latter is bound to Hp.

### IsdH binds HbHp but does not support iron acquisition by the Isd system

To assess the role of IsdH in sustaining *S. aureus* growth by mediating iron acquisition, an in-frame deletion mutant of *isdH* was constructed in the *S. aureus* wild-type (WT) background, yielding Δ*isdH*, and an *isdH* deletion was introduced into *S. aureus* Δ*isdB* strain previously generated in Hijazi et al.(Hijazi *et al*, 2026), resulting in the double Δ*isdB*Δ*isdH* strain. Iron-dependent regulation of *isdB* and *isdH* was confirmed by Fur titration assay (FURTA) (Stojiljkovic *et al*, 1994), showing that both promoters are Fur-regulated (**Figure S8**), consistent with previous reports (Dryla *et al*, 2003; Torres *et al*, 2006).

The contribution of the two hemophores was then evaluated by growing the *S. aureus* WT, Δ*isdB*, Δ*isdH* and Δ*isdB*Δ*isdH* strains under iron-restricted conditions(Hijazi *et al*, 2026) supplemented with either Hb or HbHp (120 nM) as sole iron sources (**Figure 4a,b**). As expected(Hijazi *et al*, 2026; Torres *et al*, 2006), whereas WT and Δ*isdH* grew comparably, Δ*isdB* and the Δ*isdB*Δ*isdH* mutants failed to grow in the presence of Hb, confirming the dominant role of IsdB in heme acquisition from free Hb (**Figure 4a**). In contrast, none of the strains – including *S. aureus* WT – grew when HbHp was provided (**Figure 4b**), indicating that IsdH binding to HbHp does not account for iron utilization under these conditions. As control, the addition of an excess of inorganic iron was very potent in supporting the growth of all *S. aureus* strains (**Figure 4c**).

**Figure 4.**
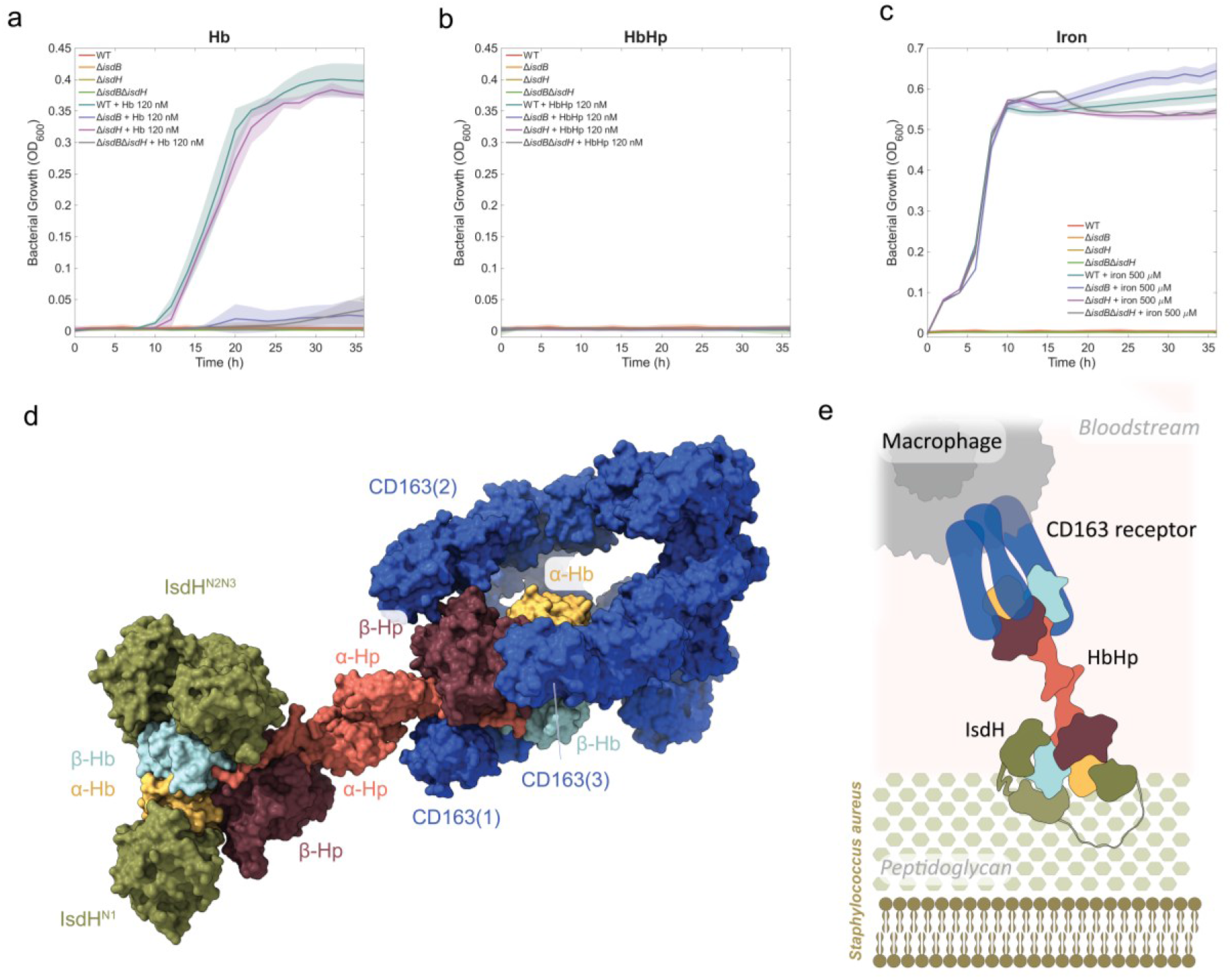
IsdH binding to the HbHp does not support iron acquisition but may promote interaction with host receptors. (**a**) Growth of *S. aureus* strains using Hb (120 nM) as the sole iron source. The WT strain and the Δ*isdH* mutant display comparable growth, whereas the Δ*isdB* mutant and the Δ*isdB*Δ*isdH* double mutant fail to grow, confirming the dominant role of IsdB in heme acquisition from free Hb. (**b**) Growth assays performed with the HbHp (120 nM Hb, 60 nM Hp) as the sole iron source. None of the strains tested, including the WT, display appreciable growth, indicating that HbHp cannot be efficiently utilized as an iron reservoir by *S. aureus*. (**c**) Growth assays performed under iron-replete conditions (500 μM FeCl_3_). (**d-e**) IsdH binding to the HbHp complex mediates binding to CD163 macrophage receptors. (**d**) Synthetic model of a tripartite IsdH:HbHp:CD163 assembly. The model was obtained by superimposing PDBs 29JK and 8XMP using the common HbHp core as structural reference. (**e**) Schematic representation of the CD163-mediated macrophage interaction with the IsdH:HbHp complex.

These results contrast with previous observations at higher HbHp concentrations (500 nM), where growth was reported for both WT and Δ*isdH* (Dryla *et al*, 2003). We propose that at elevated ligand concentrations, non-specific interactions may release sufficient heme to sustain IsdB-dependent (or alternative) uptake, thereby masking any contribution of IsdH. Under the more stringent conditions used here (120 nM HbHp), this effect is not observed, revealing the inability of *S. aureus* to efficiently exploit HbHp as an iron source. More generally, iron starvation and the consequent relief of Fur repression are recognized as global host-entry signals that activate virulence-associated pathways rather than solely iron acquisition mechanisms. Accordingly, although IsdH is induced under iron limitation, its role may extend beyond iron uptake.

Complementary bioinformatic evidence further supports a distinct functional specialization for IsdH in *S. aureus*. Comparative phylogenetic and genomic context analyses of IsdH homologues across staphylococci (**Figure S9**) reveal that only a limited number of species encode both IsdH and IsdB sequences. In these microorganisms, *isdB* co-localizes within the canonical *isd* locus alongside other genes encoding components of the heme acquisition machinery, whereas IsdH is encoded in a distinct genomic region. This separation may suggest that, unlike IsdB, whose expression is tightly coordinated within the *isd* operon and directly linked to heme uptake, IsdH might likely be subjected to different regulatory constraints. Consistently, in species lacking IsdB, IsdH homologues are often embedded within the *isd* locus, supporting a role in sustaining bacterial growth through heme acquisition.

Taken together, these data suggest that, in *S. aureus*, IsdH binding to HbHp is not associated with iron acquisition through the Isd system, although its expression is regulated by Fur. Rather, the stable interaction observed by cryo-EM is consistent with a role in mediating strong bacterial association with the HbHp complex.

### A structural model for a tripartite IsdH:HbHp:CD163 complex

CD163 is the primary macrophage receptor responsible for the clearance of HbHp via endocytosis (Graversen *et al*, 2002). Previous studies have shown that adding IsdH in solution inhibits CD163-mediated internalization of HbHp, suggesting that binding to the hemophore interferes with receptor recognition^28^. Our structural data provide a mechanistic framework for this observation. Cryo-EM analysis revealess the formation of stable 2IsdH:2Hb_dim_:Hp_dim_ complex (**Figure 1c**), which remains tightly associated without enabling iron acquisition from heme. In this configuration, IsdH engages the Hb dimer in a manner that shields interaction surfaces otherwise accessible to host receptors. Mapping the CD163-binding site onto the same Hb protomer reveals a substantial overlap with the IsdH interface, indicating that simultaneous binding of IsdH and CD163 is likely sterically hampered (**Figure S10**). Together, these findings support a model in which IsdH, when present in solution, binds to HbHp and occludes the CD163 recognition surface, thereby preventing receptor engagement and impairing macrophage-mediated clearance.

However, the situation is expected to differ when IsdH is anchored to the bacterial cell wall. In this physiological context, IsdH is spatially constrained and presented at the bacterial surface. Our structural model indicates that IsdH binding to one protomer of the Hb dimer leaves the second protomer accessible for CD163 engagement. This arrangement appears sterically compatible with the formation of a tripartite complex, in which HbHp bridges IsdH on the bacterial surface and CD163 on macrophages (**Figure 4d**). Such a configuration would effectively tether macrophages to the bacterial surface through the HbHp complex (**Figure 4e**).

## Discussion

The cryo-EM reconstruction of the complex between the full-length, physiologically relevant construct of IsdH and the HbHp complex provides a structural framework for understanding the function of this staphylococcal hemophore. The structure reveals how IsdH recognizes Hb within the HbHp complex and uncovers an unexpected modulation of subunit selectivity that depends on the presence of Hp. In addition, this work investigates the intrinsic conformational flexibility of Hp, providing insight into structure and full dynamics of this human glycoprotein unhindered by constraints imposed by a crystallographic lattice. These structural insights are supported by functional analysis, together providing a more comprehensive view of the biological role of IsdH.

The classical view of IsdH function centers on heme iron acquisition. However, the structural and biochemical analyses presented here suggest that IsdH may not primarily participate in iron scavenging but instead may contribute to other functions such as modulation of host immune responses. Indeed, IsdH forms a stable complex with HbHp but does not support bacterial growth when HbHp is provided as iron source (**Figure 4b**). This finding suggests that HbHp could instead physically bridge bacteria and macrophages through the scavenger receptor CD163.

CD163 is expressed primarily by macrophages, particularly those with an M2-like phenotype, and mediates clearance of HbHp(Murray, 2017). Macrophages are increasingly recognized not only as immune effector cells but also as potential intracellular reservoirs for *S. aureus* (Pidwill *et al*, 2021). Although traditionally considered an extracellular pathogen, *S. aureus* can survive within macrophages, especially when these cells adopt an M2-like phenotype (Lathram & Radka, 2025), potentially acting as “Trojan horses” that promote bacterial persistence and dissemination.

Within this framework, the ability of IsdH to bind HbHp at the bacterial surface, supported by NEAT1 as an additional Hb-binding domain, may promote recruitment of CD163-positive macrophages to the site of infection. The resulting tripartite interaction could bring macrophages into close proximity with the bacterial surface, potentially facilitating bacterial uptake or interactions with host cells permissive for intracellular survival.

At the same time, this interaction may be interpreted from the host perspective as a surveillance mechanism. Given that *S. aureus* predominantly exists as a commensal under conditions of immune tolerance, engagement of the HbHp complex by IsdH may act as a molecular cue associated with active iron acquisition and bacterial proliferation.

In this context, CD163-positive macrophages may exploit the IsdH:HbHp complex to selectively recognize and target *S. aureus* transitioning toward a more virulent phenotype, thereby contributing to the re-establishment of host control.

This proposed role is also consistent with the physiological context in which IsdH is expressed. Under iron-limited conditions, *S. aureus* activates a coordinated response that includes secretion of hemolysins and upregulation of the Isd system. Hemolysins promote erythrocyte lysis, releasing Hb that is rapidly captured by circulating Hp to form HbHp complexes. While the Isd system generally exploits Hb-derived heme as an iron source, IsdH displays a distinct expression profile: under iron starvation its expression increases mainly during the stationary phase of bacterial growth (Visai *et al*, 2009). This delayed expression suggests that IsdH may contribute less to rapid iron acquisition during early growth and more to bacterial persistence during later stages of infection. In this context, HbHp binding may provide indirect advantages by modulating the local immune environment rather than directly supplying heme iron.

A particularly interesting case is the epidemic USA300 strain of *S. aureus*, which exhibits altered regulation of IsdH expression due to loss of Fur-mediated repression, resulting in constitutive IsdH production even under iron-replete conditions (Poudel *et al*, 2025). Notably, USA300 also shows enhanced survival within macrophages (Tranchemontagne *et al*, 2016). Although a direct mechanistic link remains to be established, our data support the idea that altered IsdH expression could contribute to macrophage interactions that favor intracellular persistence.

Future studies will be required to test the proposed tripartite IsdH:HbHp:CD163 complex hypothesis and determine whether HbHp capture influences macrophage recruitment or polarization during infection. Investigating these mechanisms may reveal new insights into how *S. aureus* manipulates host immune responses and uncovers previously unrecognized links between iron acquisition systems and immune evasion. More broadly, the structural and functional observations reported here refine our understanding of the respective roles of the staphylococcal hemophores IsdH and IsdB, suggesting that these proteins may not be functionally redundant but instead contribute to pathogen-host interaction through distinct mechanisms.

## Material and methods

### Protein expression and purification

Expression and purification of StrepTag®II-IsdH were carried out as described in Supplementary Methods. Human hemoglobin A (HbA) was purified from outdated blood from non-smoking donors following established protocols (Viappiani *et al*, 2014). Red blood cells were lysed under hypotonic conditions, and HbA was purified by cation-exchange chromatography on CM-Sephadex C-50 using a linear pH gradient. Purified HbA was dialyzed, aliquoted, flash-frozen, and stored at ™80 °C. Methemoglobin (MetHb) was prepared by oxidation of oxyHb with potassium ferricyanide and desalted by gel filtration. Commercial haptoglobin (Hp1-1) was purchased by Athens Research & Technology Inc., and protein purity was verified by SDS-PAGE under both reducing and non-reducing conditions.

### Size-exclusion chromatography coupled with light scattering (SEC-LS)

Size-exclusion chromatography coupled with light scattering (SEC-LS) was performed using an Agilent 1260 Infinity HPLC system (Agilent Technologies) equipped with a diode array detector (1260 Infinity II WR DAD; 190–950 nm), a dual-angle LS detector (1260 Infinity MDS), and a refractive index detector (1260 Infinity II RID). Separations were carried out on a Zenix-C SEC 300 column (4.6 × 150 mm, 2.5 mL; Sepax Technologies). Column calibration was performed using a mixture of standard proteins (1 mg/mL each): ferritin (440 kDa), alcohol dehydrogenase (140 kDa), conalbumin (75 kDa), and myoglobin (17 kDa). LS Detector calibration was conducted using bovine serum albumin (BSA) at an approximate concentration of 10 g/L.

### Cryo–EM sample preparation and data acquisition

To investigate the molecular architecture of the ternary complex formed by IsdH and HbHp, cryo-EM grids were prepared following an optimized protocol previously described(De Bei *et al*, 2022). The complex was prepared at a 1:2:1 molar ratio of IsdH^N1N2N3^, metHb (heme basis), and Hp1-1. Samples were prepared at a final concentration of 6 g/L in 10 mM HEPES buffer (pH 7.2), selected to minimize background electron density, and supplemented with 3.5 mM CaCl_2_. To mitigate preferred particle orientation in vitreous ice, the zwitterionic detergent 3-([3-Cholamidopropyl]dimethylammonio)-2-hydroxy-1-propanesulfonate (CHAPSO) was added immediately prior to vitrification to a final concentration of 8 mM.

Three microliters of sample were applied to glow-discharged 300-mesh R1.2/1.3 QUANTIFOIL® grids, blotted for 3 s, and plunge-frozen in liquid ethane using a Vitrobot Mark IV (Thermo Fisher Scientific) operated at 4 °C and 99% humidity. Initial grid screening was performed on a Talos Arctica transmission electron microscope operating at 200 kV and equipped with a Falcon 3 linear detector (School of biological sciences Cryo-EM Facility, University of Cambridge).

High-resolution data collection was carried out on a Titan Krios transmission electron microscope (Thermo Fisher Scientific) operating at 300 kV and equipped with a Falcon 4i direct electron detector and a SelectrisX energy filter (Thermo Fisher Scientific) set to a 10 eV slit width (School of Biological Sciences Cryo-EM Facility, University of Cambridge). Movies were acquired using EPU software (Thermo Fisher Scientific) at a calibrated magnification of ×165,000, corresponding to a physical pixel size of 0.729 Å. Each micrograph consisted of 50 frames collected over 4.39 s, resulting in a total dose rate of 12.24 e^−^/Å^2^/s. Data were collected with a nominal defocus range of −0.6 to −1.8 µm. In total, 6,324 micrographs were acquired for the IsdH:HbHp complex.

### Cryo-EM single-particle analysis and flexibility assessment

Single-particle analysis (SPA) was performed using RELION 5.0(Scheres, 2012) and cryoSPARC (Punjani *et al*, 2017). Raw movies were motion-corrected with MotionCor2 (Zheng *et al*, 2017), and CTF parameters were estimated using CTFFIND 4.1 (Rohou & Grigorieff, 2015). Particles were picked with the TOPAZ neural network(Bepler *et al*, 2019), trained, when necessary, with 2D classification projections to improve selection. Multiple rounds of 2D and 3D classification were used to solve compositional heterogeneity and isolate high-quality particle subsets. 3D refinement was followed by CTF refinement and Bayesian polishing to generate high-resolution reconstructions.

Homogeneous particle subsets were imported into SCIPION (De La Rosa-Trevín *et al*, 2016) for detailed analysis of conformational flexibility. Continuous structural dynamics were examined using 3DFlex (Punjani & Fleet, 2023) and Multibody Refinement (Nakane & Scheres, 2021).

Atomic models of the IsdH:HbHp complex were generated using AlphaFold3 (Abramson *et al*, 2024) and flexibly fitted into density maps with Namdinator (Kidmose *et al*, 2019). Ambiguous or flexible regions were manually adjusted in ISOLDE (Croll, 2018) to achieve optimal agreement with the cryo-EM density.

### Immunoassay for determination of hemophore–HbHp binding affinity

Binding affinities of IsdH constructs for the HbHp complex were determined using a Strep-Tactin–based immunoassay adapted from previous work(Buoli Comani *et al*, 2025), in which immobilized StrepTag-fused hemophores were incubated with increasing concentrations of HbHp and binding was quantified using an HRP-conjugated anti-Hb antibody. Dissociation constants were obtained by fitting the data to a one-site binding model (see Supplementary Methods for details).

### Bacterial strains and growth conditions

Growth assays were performed using *S. aureus* Newman WT and isogenic Δ*isdB*, Δ*isdH* and Δ*isdB*Δ*isdH* mutants generated by allelic replacement. Bacteria were cultured under iron-limited conditions in the presence or absence of Hb or HbHp, and growth was monitored by OD_600_ measurements (see Supplementary Methods for strain construction and experimental details)

## Acknowledgments

This project was funded by “PRIN-2020 – Defeat antimicrobial resistance through iron starvation in *Staphylococcus aureus* (ERASE)” (Grant 2020AE3LTA) to SB and EF. This research benefits from the High-Performance Computing facility of the University of Parma, Italy (https://www.hpc.unipr.it). This work has been carried out in the frame of the ALIFAR project, funded by the Italian Ministry of University through the program ‘Dipartimenti di Eccellenza 2023-2027’. VBC received cryo-EM training through Wellcome/MRC funded program (218785/Z/19/Z). ODB received cryo-EM training through Wellcome Trust and MRC funded Cryo-EM Training Program (Advanced Single Particle Cryo-EM Course, 2025).

